# Block aligner: fast and flexible pairwise sequence alignment with SIMD-accelerated adaptive blocks

**DOI:** 10.1101/2021.11.08.467651

**Authors:** Daniel Liu, Martin Steinegger

## Abstract

**Background:** The Smith-Waterman-Gotoh alignment algorithm is the most popular method for comparing biological sequences. Recently, Single Instruction Multiple Data methods have been used to speed up alignment. However, these algorithms have limitations like being optimized for specific scoring schemes, cannot handle large gaps, or require quadratic time computation.

**Results:** We propose a new algorithm called block aligner for aligning nucleotide and protein sequences. It greedily shifts and grows a block of computed scores to span large gaps within the aligned sequences. This greedy approach is able to only compute a fraction of the DP matrix. In exchange for these features, there is no guarantee that the computed scores are accurate compared to full DP. However, in our experiments, we show that block aligner performs accurately on various realistic datasets, and it is up to 9 times faster than the popular Farrar’s algorithm for protein global alignments.

**Conclusions:** Our algorithm has applications in computing global alignments and *X*-drop alignments on proteins and long reads. It is available as a Rust library at https://github.com/Daniel-Liu-c0deb0t/block-aligner.

## Background

Efficient comparison of biological sequences is an incredibly important problem due to its prevalence in many bioinformatics workflows and the ever-increasing scale of experiments. For example, protein sequences are aligned in large-scale database searches [1, 2]. In sequencing experiments, millions of reads are compared to each other or aligned to references [3, 4, 5]. Currently, the most popular method for comparing biological sequences is computing optimal pairwise alignment (weighted edit distance) with the dynamic programming Smith-Waterman-Gotoh algorithm [6] (and its global variant, the Needleman-Wunsch algorithm [7]).

The Smith-Waterman-Gotoh algorithm and its variants compute the alignment score and path by rewarding matches and penalizing insertions, deletions, and substitutions between two sequences. To obtain biologically relevant results, an affine gap model is often used, where starting a new gap (insertion or deletion) and continuing an existing gap have different costs [6]. There are many variants of this algorithm, though in this paper we mainly focus on global alignment (end-to-end alignment of two sequences) and *X*-drop alignment (align some prefix of two sequences until the scores drop by *X* below the maximum score, terminating alignment early for dissimilar sequences). Additionally, we aim to address both nucleotide and protein sequence alignment in this paper. For protein alignment, there is the added complexity of using 20 × 20 amino acid substitution matrices (*e*.*g*., BLOSUM62) [8], instead of 4 × 4 nucleotide substitution matrices. However, despite the versatility and usefulness of the Smith-Waterman-Gotoh algorithm and its variants, they typically scale quadratically with the lengths of the input sequences, making them prohibitively slow on long sequences.

Since this alignment step is so critical, countless optimizations have been introduced in the past. Recent advances in alignment algorithms have made use of Single Instruction Multiple Data (SIMD) instructions in most modern CPUs to speed up the DP computation. For example, the x86 AVX2 vector extensions allow many basic arithmetic operations to operate on multiple lanes in a 256-bit vector register in parallel. Here, we briefly review the most popular previous techniques for accelerating sequence alignment with SIMD. In Figure 1, we show the most popular methods for tiling SIMD vectors.

**Figure 1:**
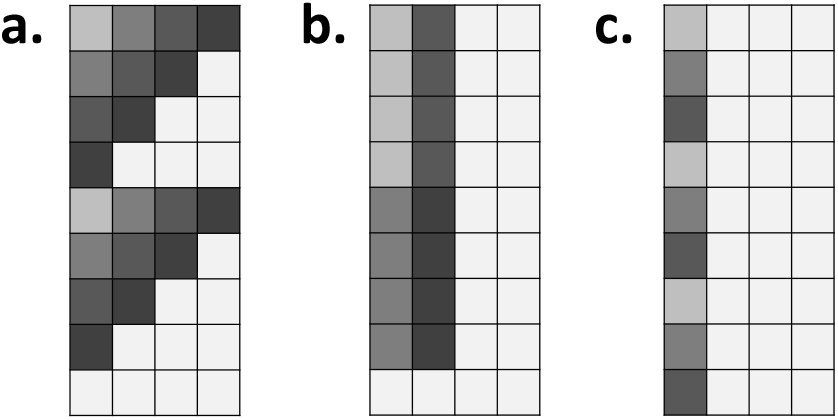
Vectorization strategies. Cells in the DP matrix with the same color can be computed in parallel with a single SIMD vector. (a) Vectors are laid out anti-diagonally. (b) Vectors are tiled vertically. Dependencies between adjacent cells within the same vector are typically resolved through prefix scan. (c) Striped layout where vectors hold noncontiguous cells within each column.

### Bit parallel methods

Myer’s algorithm [9] and its variants were designed for computing edit distance, where two strings are aligned by penalizing mismatches, insertions, and deletions with unit costs (Levenshtein distance), using general purpose registers as bit vectors. This simple form allows differences between DP cells to be computed with basic bit operations. This method has been extended to use wider SIMD registers and cells can be pruned based on an edit threshold for greater speedups in edlib [10]. A generalization to handle integers scores was proposed in BitPAl [11]. However, despite the high amount of parallelism achieved with bit vectors, these methods are cannot handle affine gaps and complex scoring matrices.

### Anti-diagonal methods

Since cells in the same antidiagonal in the Smith-Waterman-Gotoh DP matrix do not depend on each other, anti-diagonal cells can be computed in parallel [12] with SIMD vectors. In practice, this is done by tiling an anti-diagonal vector across the entire DP matrix. Although this avoids any over-head in resolving dependencies between anti-diagonal cells, this also makes looking up scores in large protein scoring matrices for each lane in a SIMD vector slow.

### Static banding

If the number of insertions or deletions is bounded, then one way to speed up the Smith-Waterman-Gotoh algorithm is only computing scores in a (static) band of a fixed width around the diagonal of the DP matrix [4]. SIMD vectors are typically tiled anti-diagonally within this band to avoid resolving dependencies within the vectors. However, this method requires the band width to be known in advance, and allowing more insertions or deletions drastically reduces the speed of the algorithm.

### Adaptive banding

To achieve greater speedups over static banding, while risking reducing accuracy, the anti-diagonal band can be made adaptive by greedily shifting it down and right based on the location of the best score [13]. Compared to the static banding approach that only computes DP cells around the diagonal, adaptive banding allows much smaller band widths to be used by allowing the band to stray from the diagonal. To handle large scores that accumulate in long sequences without using wider SIMD lanes, the Suzuki-Kasahara difference recurrence algorithm [14] was proposed to only save differences between DP cells. Since differences between cells are small, this allows for maximum parallelism with SIMD vectors. Conceptually, this algorithm is similar to Myer’s edit distance algorithm, but it is extended to support affine gap scores with SIMD vectors. One major drawback of the adaptive banding approach is that the greedy algorithm for shifting the band cannot easily handle large gaps, causing it to return a suboptimal alignment score. Additionally, the anti-diagonal bands are not amenable to aligning protein sequences.

### BLAST X-drop algorithm

If an user-defined *X*-drop threshold is available, it is possible to prune DP cells that drop by more than *X* below the max score so far. This method can be SIMD vectorized by computing rows in the DP matrix and pruning vectors if they meet the *X*-drop criteria, until there are no vectors left [13]. Like banded methods, if the *X*-drop threshold is large, then the algorithm will be very slow. Also, it is difficult to obtain global alignments with this algorithm.

### Striped methods

Farrar’s algorithm [15] is a SIMD-accelerated algorithm for computing the entire DP matrix. In contrast to banded approaches, each SIMD vector is used to hold scores from strided, noncontiguous cells in a column in the DP matrix. It requires an extra loop for lazily fixing up vertical dependencies. A similar algorithm is Daily’s prefix scan variant [16], which also uses the striped pattern for storing cells. Since these striped methods compute the full DP matrix, they are suitable for both global and local alignment [17]. However, these methods require preprocessing the input sequences in order to make efficient retrieval of noncontiguous characters of the sequences possible.

### Wavefront Alignment

Recently, the Wavefront Alignment method [18] has been proposed that excels on similar sequences. This method is able to iteratively increase the positive score threshold and expand the computed region of cells by only storing diagonal lengths that represent the computed wavefront. This method is able to identify exactly matching regions to rapidly align sequences with high sequence identity. However, it is limited to scoring schemes where gaps and mismatches increase the score and the goal is to minimize the overall score.

### Intersequence parallelization

Although we focus on *intrasequence* parallelization techniques for pairwise sequence alignment in this paper, there also exists methods for exploiting *intersequence* parallelization by aligning multiple sequences against a single sequence [19]. This is typically done by having each lane in a SIMD vector hold the DP score for a different pair of sequences. Although this is simpler than intrasequence parallelization since there are no dependencies between lanes, the benefit of this approach only appears when there are many input sequences to be aligned.

Other algorithms exist for GPUs (*e*.*g*., Logan [20]) and other special devices. Aligning a sequence against a graph (*e*.*g*., abPOA [21]), or even aligning raw Nanopore signals [22]. However, they are out of the scope of this paper as we focus on pairwise sequence alignment on the CPU.

Despite these many recent advances in improving pairwise sequence alignment, many methods cannot accurately and efficiently handle complex protein scoring schemes, long alignment gaps, global alignment, and *X*-drop alignment. In this paper, we propose a new SIMD-accelerated algorithm for pairwise alignment of protein and nucleotide sequences, called “block aligner”. It draws ideas from adaptive banding [13] to allow a small block size to be used compared to commonly used static banding approaches [12]. Also, the block size is able to grow in order to span larger gaps that would mislead a naive adaptive banding approach. Our unique approach using blocks tiled with horizontal and vertical SIMD vectors makes handling complex scoring matrices (*e*.*g*. for amino acids) possible. In exchange for these features, block aligner provides no theoretical guarantees that the correct results will be produced, unlike computing the entire DP matrix [6, 15, 16] or other methods like Wavefront Alignment algorithm [18]. However, in our experiments, we show that block aligner performs well in practice on both synthetic and real protein and sequencing read datasets. For example, it reaches up to 9 times faster than Farrar’s algorithm [15] on protein global alignment, while maintaining high alignment accuracy.

## Methods

### Smith-Waterman-Gotoh Alignment

Here, we introduce the Smith-Waterman-Gotoh algorithm [6], along with its global variant, the Needleman-Wunsch algorithm [7] in detail. They are dynamic programming techniques that compute the optimal alignment of two sequences *q* and *r* (of length |*q*| and |*r*|, respectively) by computing scores in a *O*(|*q*||*r*|) matrix along with the transition directions (trace). After this matrix is computed, we can easily find the sequence of substitution, insertion, or deletion edits to transform one sequence to the other, along with the score of such an alignment. Affine gap (insertion or deletion) penalties of the form *G*_*ext*_(*g −* 1) + *G*_*open*_ are typically used, where *G*_*ext*_ and *G*_*open*_ are gap scoring parameters and *g* is the number of gaps. For our cases of *X*-drop and global alignment, the DP recurrence is the same:

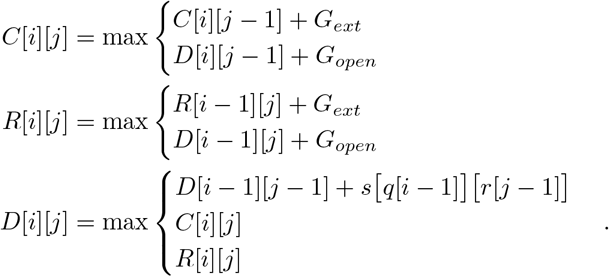

Note that we use *D, R*, and *C* (*R* and *C* stand for “row” and “column”) in our notation instead of *H, E, F* from the original Farrar paper [15].

For the boundary conditions, we have

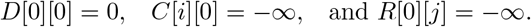

In global alignment, the optimal score is *D*[|*q*|][|*r*|]. In *X*-drop alignment, it is the maximum computed *D* score. The *X* threshold is useful in terminating alignments early when the scores drop by *X* below the maximum computed score so far.

In this paper, we assume that the CPU supports SIMD vectors with *L* 16-bit lanes that can be operated on with some basic operation in parallel. These lanes would typically hold the DP scores as 16-bit integers.

### Overview of Block Aligner

In contrast to popular diagonal banding approaches, block aligner makes use of a novel block approach. This approach makes use of vertical or horizontal vectors that are laid out to form a square block. Like adaptive banding [13], this block is greedily shifted right or down based on where the score seems to be the largest. Compared to static banding approaches, such an adaptive approach allows a small band or block size to be used even if the optimal scores stray far from the diagonal (see the Sequencing Reads Benchmark section).

A major limitation of previous adaptive banding approaches is that at the start of a gap, it is hard to differentiate whether the gap is in one sequence or the other, or whether there is just a region of many mismatches with a small band. In these cases, the band must span the length of the gap in order for the direction to shift to be accurately determined. In block aligner, this issue is solved by allowing the block size to grow. Based on a heuristic similar to *X*-drop, block aligner will dynamically recompute some regions in the DP matrix and double the block size to handle low sequence identity segments in the sequences. Unlike the BLAST *X*-drop algorithm that prunes cells only when they fail the *X*-drop test, block aligner is more optimistic: a smaller block size is assumed to start with, and it only grows when necessary. This shares some similarities with the Wavefront Alignment algorithm [18], where higher sequence similarity equates to less computed DP cells.

### The Block Aligner Algorithm

We show pseudocode for our algorithm in Algorithm 1, which we use to guide our explanation of block aligner. We also give a diagram of block aligner in Figure 2. To help visualize the cells computed by block aligner, we draw the computed rectangular regions for two pairs of real proteins in Figure 3.

**Figure 2:**
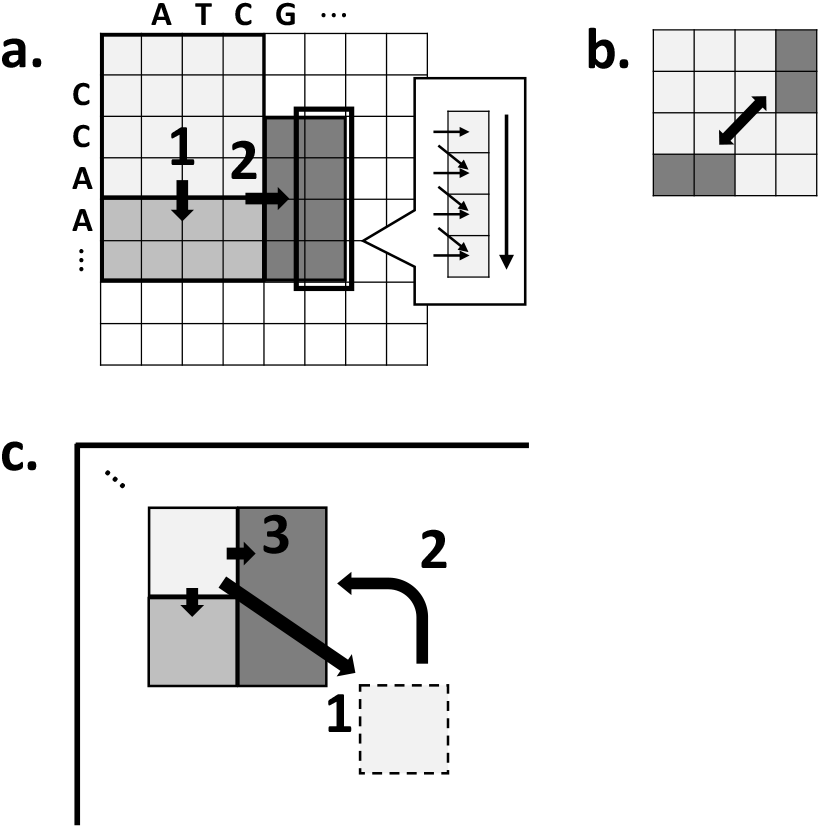
Notable aspects of the block aligner algorithm. (a) 1. The block is shifted down by a step size of *S* = 2, and only new cell scores are computed. 2. The block is shifted right. Each rectangular region is computed by tiling horizontal or vertical SIMD vectors. Prefix scan is used to resolve cell dependencies across a vector. (b) The sum of the *S* cells in the bottom and right borders of a block are compared to determine the direction to shift. (c) 1. The block is shifted down and right. 2. When the *Y* -drop criteria is met, some computed cells are discarded to return to a previous checkpoint state. The block doubles in size and alignment restarts.

**Figure 3:**
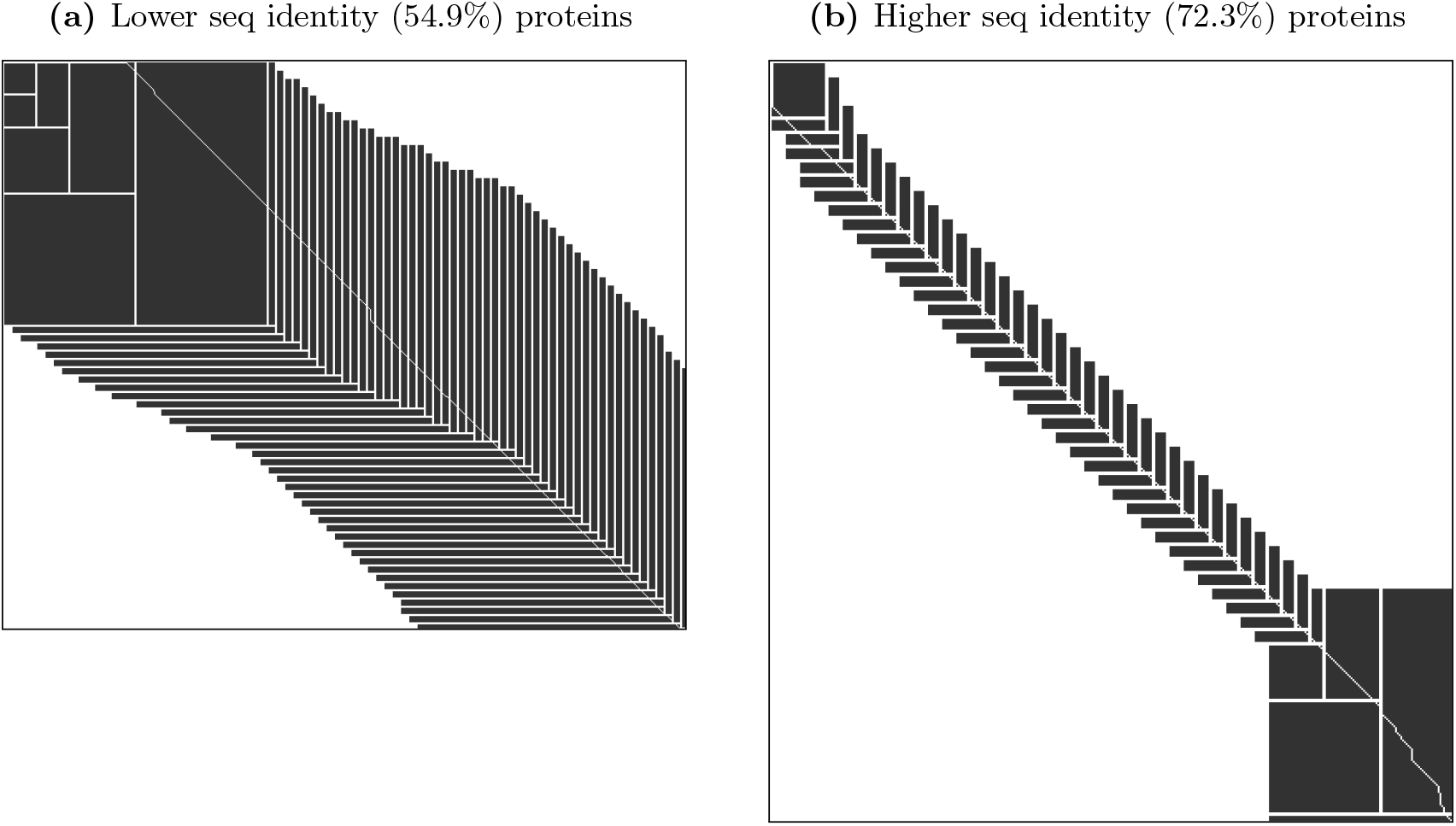
Block aligner visualized. All regions computed by block aligner when aligning pairs of protein sequences from the Uniclust30 dataset are shaded. The thin white line stretching from the top left corner to the bottom right corner is the traceback path of the optimal alignment.

#### Computing Scores

Block aligner relies on the Compute Rect subroutine in Algorithm 2 to efficiently compute the scores for certain rectangular regions of the DP matrix that has the top left corner at (*r, c*) and with dimensions (*w, h*). When shifting right or down, a rectangular region to the right or below the current block is computed. When **A** growing a block, two rectangular regions must be computed, at the bottom and to the right of the current block, in order to piece together a larger block. It is wasteful to recompute the scores for the whole block, so we only compute the new regions that are exposed after a shifting or growing. Block aligner defaults to using a step size of *S* = 8 for shifting, which is typically smaller than the size of the vector *L* (*L* = 16 with AVX). This means that the SIMD vectors need to be tiled vertically for right shift and horizontally for down shift. The same Compute Rect function can be easily reused for right and down shifts by swapping *R* and *C*.

Compared to naively implementing the Smith-Waterman-Gotoh algorithm, our algorithm has many differences. When tiling vectors vertically and horizontally, dependencies between lanes within the SIMD vectors (computing the *R*′ array) are resolved with prefix scan, an *O*(log *L*) time operation. This has been explored previously in [13, 21] on CPU and in [23] on GPU. However, our implementation has a couple of important optimizations: for 256-bit AVX vectors, there is a performance penalty for moving values between its lower and higher 128-bit lanes. Therefore, we avoid this by performing prefix scan on the 128-bit lanes separately using the Kogge-Stone method [24], then doing one lane-crossing shuffle to resolve the dependency from the between the upper and lower halves of SIMD vector like in the Sklansky method [25]. Additionally, we reorder the dependency of *R*[*i* + *i*′][*j*] on the *R*[*i −* 1][*j*] value of the previous vector to be after the prefix scan step. This allows some instruction-level parallelism to be exploited, since prefix scans for computed *R*′ from multiple iterations of the loop are independent. Similar ideas have been explored in the context of GPU prefix scans [26], but we extend them to sequence alignment with CPU SIMD instructions.

##### Algorithm 1 Pseudocode for the block aligner algorithm.

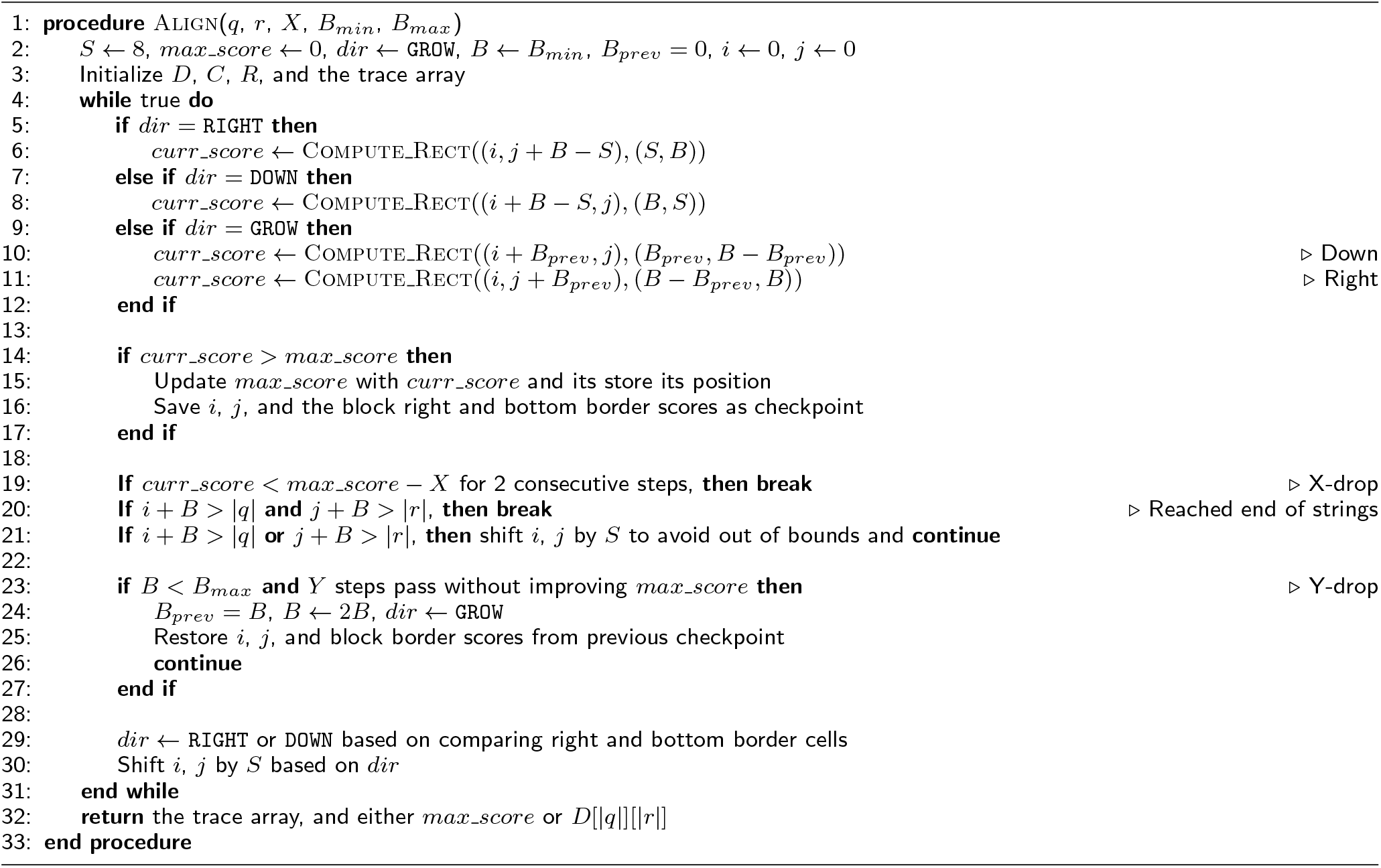

##### Algorithm 2 Pseudocode for computing cells in a region of the DP matrix.

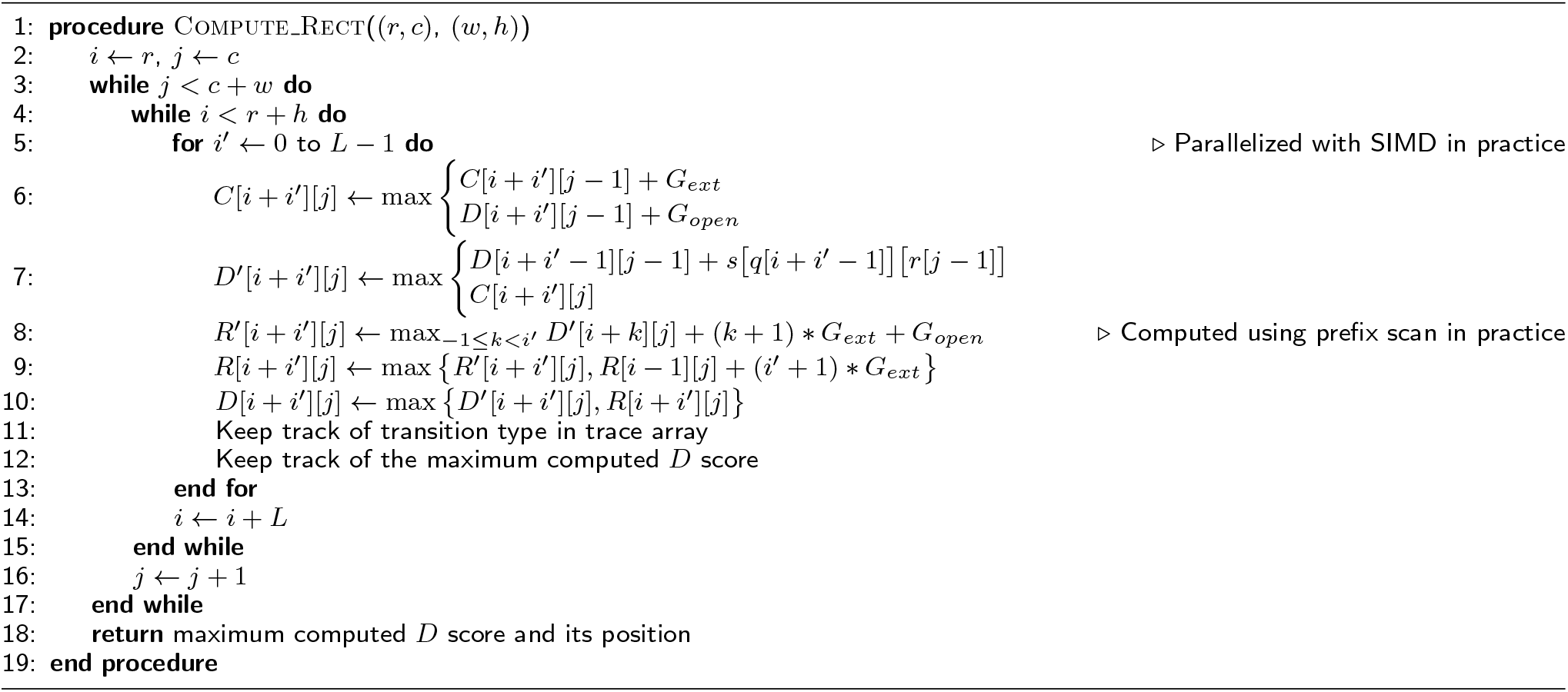

We do not tile vectors anti-diagonally [12] because it makes looking up scores for multiple lanes in the SIMD vector difficult. Loading scores for a horizontal or vertical vector in the DP matrix is much easier: a single row of the scoring matrix is loaded, and the scores for each cell in the SIMD vector can be determined through fast vector shuffles (*e*.*g*., a combination of `_mm256_shuffle_epi8` and `_mm256_blendv_epi8` to lookup a 32 element row in the score matrix with a 5 bit index). This is vital to support protein scoring matrices of size 20 × 20.

We do not use the striped profile in Farrar’s algorithm [15] and Daily’s prefix scan variant [16] because they require precomputed striped profiles for the query sequence, which is not possible to obtain when the computed block shifts dynamically.

In contrast to adaptive banding, block aligner accurately identifies the maximum score from anywhere in the block, instead of only examining the center of the band.

#### Offsets

Although not shown in the pseudocode, we use 16-bit lanes to represent relative scores and 32-bit offsets for each block. Since we cannot use the difference recurrence method [14] along with prefix scans, and amino acid scoring matrices typically have large scores, we were unable to use narrow 8-bit lanes for more parallelism. Also, our approach is more similar to that of [27], where a band has a single scalar offset. We choose the maximum score from the previous step as the current step’s offset to ensure that larger scores are accurately represented at the expense of lower scores that are often irrelevant. This offset method should accurately compute the relevant scores in most reasonable cases.

#### Determining Shift Direction

The scores on the right and bottom borders of each block are examined to determine the direction to shift the block in the next step. We compare the sum of the *S* leftmost scores in the bottom border with the sum of the *S* topmost scores in the right border:

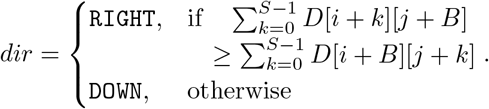

Summation allows all *S* scores in a border to contibute to the aggregate, in contrast to other operations like max. Our method differs from locating the max score in an adaptive band [27] or comparing only the bottomleft and top-right corner cells in an adaptive band [13].

*Y -drop, Block Growing, and Restoring Checkpoints* In order to handle large gaps, we save checkpoints use a heuristic to determine when there is a gap and the block size needs to grow. Every time a new maximum score is encountered, a checkpoint of the current state of the block is saved. If, after some threshold *Y* number of iterations, a new maximum score has not been reached, then the block returns to the previous checkpoint and the block size is doubled. This checkpoint restoration can happen anytime the *Y* threshold condition is met during alignment computation. The block size is allowed repeatedly grow in quick succession if a new max score is not identified after growing. Any regions that were computed after the last checkpoint are discarded and recomputed for better accuracy. This *Y* threshold is similar to the popular *X*-drop threshold, but it represents a number of iterations, which is scoring-matrix-agnostic. We fix 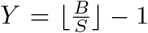, which means that the *Y* threshold is met if the block shifts approximately *B* cells away from where the last maximum score is encountered. We only increase the block size and not decrease, in order to keep the algorithm simpler and avoid potentially making mistakes when there are gaps.

#### Traceback

To obtain the exact edit operations used, we use the `movemask` instruction (*e*.*g*.,`_mm256_movemask_epi8`) to compress the DP matrix’s trace direction information and only store two bits per score in a SIMD vector, 8 times less than the 16 bits used for relative scores. This trace information is non-temporally stored into memory so it does not pollute the cache. The trace data is treated like a stack: when restoring a previous checkpoint, trace information for discarded blocks are popped off until the checkpoint is reached. To avoid eagerly allocating too much memory for trace data at the start based on the maximum block size, we estimate how much trace data needs to be stored to reach the ends of the sequences to expand the allocated memory each time the block grows. When the traceback path (*e*.*g*., CIGAR) is requested, block aligner simply follows the stored 2-bit encoded trace directions of the DP matrix starting from the end of the alignment and going back to the origin. There is room for further optimization here, like for example using the popcount approach in adaptive banding.

### Complexity

#### Time Complexity

With two sequences of length *N*, there must be less than 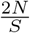 right or down steps to shift a block from the top left corner of the DP matrix to the bottom right corner. Each shift requires 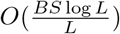 vector operations, if the block size is *B*. Assuming that the number of checkpoint restarts is insignificant compared to *N*, the overall time complexity of block aligner is 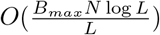. Compared to adaptive banding, block aligner has an extra factor of log *L* due to prefix scan.

#### Space Complexity

A constant number of arrays of size *B*_*max*_ are needed to keep track of the current scores in the bottom and right borders of the block, along with the scores at the previous checkpoint. Therefore, the space complexity of block aligner is *O*(*B*_*max*_). When saving traceback information, each cell that is computed must use two bits to store its previous cell. This will take *O*(*B*_*max*_*N*) extra bits of space.

#### Usage Complexity

Since block aligner is more complex than other banded approaches, we have made an effort to reduce the number of tunable parameters that may affect algorithm performance, so it works well “out of the box”. Part of this is the score-agnostic *Y* -drop parameter. We also fixed the step size to be *S* = 8 for 256-bit or longer vectors and *S* = 4 for 128-bit vectors. Therefore, the only extra parameters in addition to the typical *X*-drop threshold and (gap) scoring parameters are the block sizes *B*_*min*_ and *B*_*max*_, which must be powers of two. Since the block size can grow, block aligner is very forgiving on the choice of *B*_*min*_ and *B*_*max*_ parameters.

### Library Implementation

Block aligner is implemented as a Rust library. It makes extensive use of const generics to allow the compiler to generate efficient code specifically for certain alignment options. Currently, block aligner supports x86 AVX2 and Webassembly [28] SIMD instruction sets. C bindings are also available for block aligner.

When the block aligner library is used, the input sequences must be padded with special bytes to avoid out of bounds access with future SIMD memory loads. Also, the block size is limited to fit within a 16-bit integer to save space when storing traces. Block aligner supports protein scoring matrices (*e*.*g*., BLOSUM62 [8]), DNA scoring matrices (with undetermined “N” bases), and a simplified scoring matrix based on fixed scores for matches or mismatches when aligning arbitrary byte sequences.

Due to block aligner’s heavy usage of unsafe SIMD operations, we have carefully tested it with both manually specified test cases and randomly generated inputs. Additionally, block aligner contains many assertions to catch memory issues if it is built and ran in debug mode.

To identify speed bottlenecks and optimize them, we have profiled and carefully examined the compiled assembly code of the core block aligner implementation. This has led to many small performance wins, most commonly in cases where the generated assembly had unexpected overheads due to runtime bounds checks or other quirks of certain Rust standard library functions.

## Results

### Setup

We evaluate block aligner on both simulated and real datasets:

#### • Random DNA or proteins

We generate DNA datasets with sequences of length *N* by drawing from {A, T, C, G, N}^*N*^ with equal probability. For proteins, we use the set of all 20 amino acids. One sequence in each pair of alignable sequences is mutated by adding between 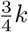 and *k* substitution, insertion, or deletion edits, with *k* being a parameter that we vary. To investigate the impact of long, consecutive gaps, we also generate data where we insert a random subsequence with length equal to 10% of the sequence length into a random position in one of the two pairs of sequences.

#### • Uniclust30

We use mmseqs2 [1] (commit 24f6b52) to find alignable sequences above a certain coverage (overlap) threshold. Two coverage thresholds are used: 80%, which is the default, and 95%, which is an easier dataset that is more globally alignable. We will refer to these Uniclust30 [2] (UniRef30 version 03 2021) datasets with different parameters as “uc30” and “uc30 0.95,” respectively. For both datasets, the proteins range from between around 20 to around 8,000 amino acids long, with an average length of around 300. There are 7,000 proteins per dataset.

#### • 25 kbp Nanopore reads

We use the 1,734 pairs of sequences derived from aligned Oxford Nanopore MinION reads of around 25,000 bases long as a long read sequencing dataset. This data is taken from [14], where it was generated by aligning reads from [29] with BWA-MEM [3]. The accession numbers are FAB45271, FAB42316, and FAB49164. This dataset does not have pairs with gaps of over 20 base pairs long. In some experiments, we only use a prefix of a certain length, compared to the “default” of using the entire sequence.

#### • Illumina and 1 kbp Nanopore reads

We also use two additional DNA read datasets. One dataset is 100,000 pairs of sequences derived from Illumina HiSeq 2000 reads of length 101, with accession number ERX069505. Another dataset is 12,477 pairs of sequences derived from Oxford Nanopore MinION reads of around 1,000 bases long from [30], with accession numbers ERR3278877 to ERR3278886. These datasets were taken from [18], where the reads were aligned with minimap2 [4].

We also evaluate block aligner against a diverse set of previous algorithms:

#### • Rust-bio [31] (version 0.33.0)

We use the scalar full DP implementation to perform global alignment.

#### • Parasail [16]

We use the saturating version of the implementation of Farrar’s striped SIMD algorithm [15] for computing the full DP matrix for global alignment. This version will attempt to align with 8-bit SIMD lanes, and only fall back to 16-bit lanes if an overflow is detected. In our experiments, Parasailors [32] (version 0.3.1), the Rust bindings for Parasail are used.

#### • Seqan [33] (version 1.4.2)

We use Seqan’s implementation of Myer’s bit vector edit distance method [9] which computes the full DP matrix for global alignment.

#### • Edlib [10] (see [34] at commit f056862)

Edlib also provides an implementation of Myer’s edit distance algorithm, but it allows the maximum number of edits to be thresholded with *k*.

#### • Static banding

We use two different static antidiagonal banding implementations: triple accel [35] (from version 0.33.0 of Rust-bio), a pure Rust library optimized for SIMD global alignment with small band widths, and the banded SIMD implementation tested in [13] (see [36] as commit dc65c0d) that only returns the max score.

#### • Adaptive banding

We test multiple implemen-tations of SIMD-accelerated adaptive banding from [14] (see [34] at commit f056862). This includes “libgaba” and “diff-raw,” which implements the difference recurrence algorithm with a band width of 32 and 8-bit lanes, “editdist,” Myer’s algorithm with an adaptive band width of 64, and “non-diff,” which is adaptive banding (band width of 32) without the difference recurrence algorithm and with wider 16-bit lanes.

When measuring the error rate of an algorithm, we typically count how many pairs of alignments are *wrong*. An alignment is wrong if its score does not match the *correct* score produced by a full DP computation. Adaptive approaches will never overestimate the score, so a wrong alignment means that the score is lower that the correct result. To measure how much a wrong score differs from the correct score, we compute the percent error, which is simply

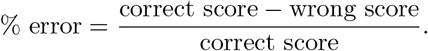

We will write *B*_*min*_-*B*_*max*_ (*e*.*g*., 32-256) to indicate the min and max block sizes. If we only specify the max block size, then the min block size defaults to 32. For scoring, we use BLOSUM62 [8] with (*G*_*open*_ = -11, *G*_*ext*_ = -1) for proteins and (match = 1, mismatch = -1, *G*_*open*_ = -2, *G*_*ext*_ = -1) for DNA sequences.

We will often refer to the sequence identity of a pair of sequences, which is defined as

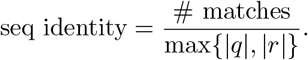

All of our benchmarks are conducted on a 2019 MacBook Pro with 2.3 GHz Intel Core i9 CPUs and 16 GB of RAM. Unless otherwise noted, our benchmarks use 256-bit AVX2 vector registers. Our experiments are fully reproducible. The code, data, and results are available in our repo: https://github.com/Daniel-Liu-c0deb0t/block-aligner.

### Random Protein and DNA Benchmark

Our first experiments are with random data, as a quick verification of block aligner. With random DNA sequences with lengths in {10^2^, 10^3^, 10^4^}, mutation rates (number of edits *k* as a percentage of the sequence length) in {10%, 20%, 50%}, and a fixed block size of 32, the error rate of block aligner is 0%. This shows that block aligner works, but random data is not very challenging and we expect the error rate on real data to be higher.

Next, we investigate whether block aligner is able to handle large gaps with its block growing heuristic, by inserting a long random subsequence per pair of sequences. Our results in Figure 4a show that block aligner’s growing heuristic is able to span extremely large gaps.

**Figure 4:**
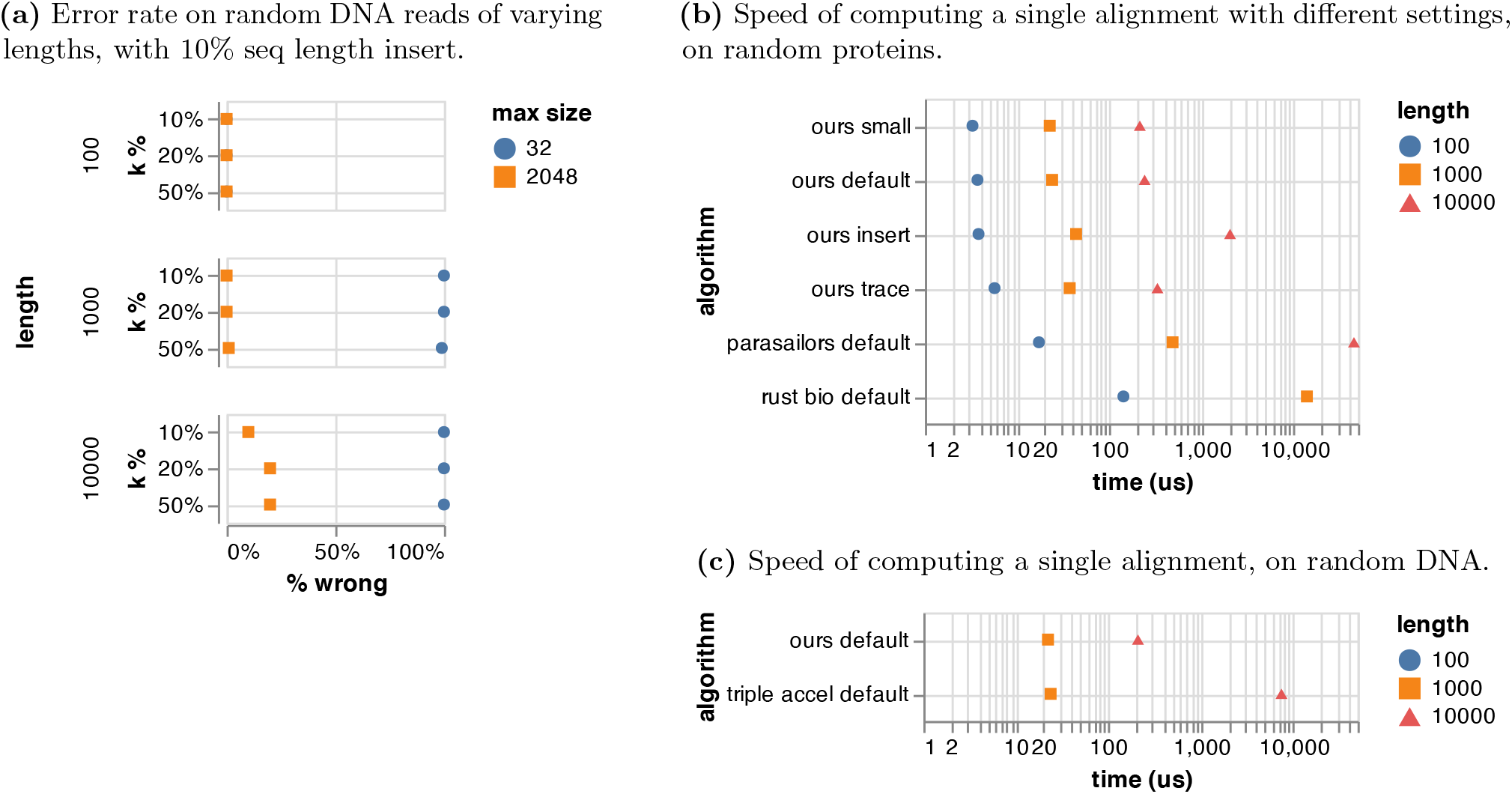
Global alignment of random DNA and protein sequences with block aligner. The default block size is 32-2048. In the benchmarks, a *k* % of 10% is used. In (a), 100 random reads were used to compute each percentage, except 10 random reads were used when the length is 10,000. In some cases in (b), other settings are used: “small” means block size is 32-32 (no growing), “insert” means that there is a 10% seq length random insert, and “trace” means that trace data is saved and the traceback is computed. In (c), triple accel uses a band width equal to 10% of the seq length.

In Figure 4, we also benchmark block aligner versus Parasail [16], Rust-bio [31], and triple accel [35]. Note that in Figure 4c, triple accel is able to use narrow 8-bit lanes when the sequence length is 1,000 and the band width is 100, and wider 16-bit lanes on longer sequence lengths. Block aligner always uses 16-bit lanes. In terms of speed, block aligner is multiple times faster than Parasail and Rust-bio, as expected. Block aligner does slow down when the block size is forced to grow to handle the large insert, but even so, it is still faster than Parasail’s full SIMD DP. Although block aligner’s traceback procedure is not highly optimized, it is not a speed bottleneck.

### Uniclust30 Protein Benchmark

To assess how block aligner performs on real protein data, we align pairs of proteins from the Uniclust30 dataset [2] and report our results in Figure 5.

**Figure 5:**
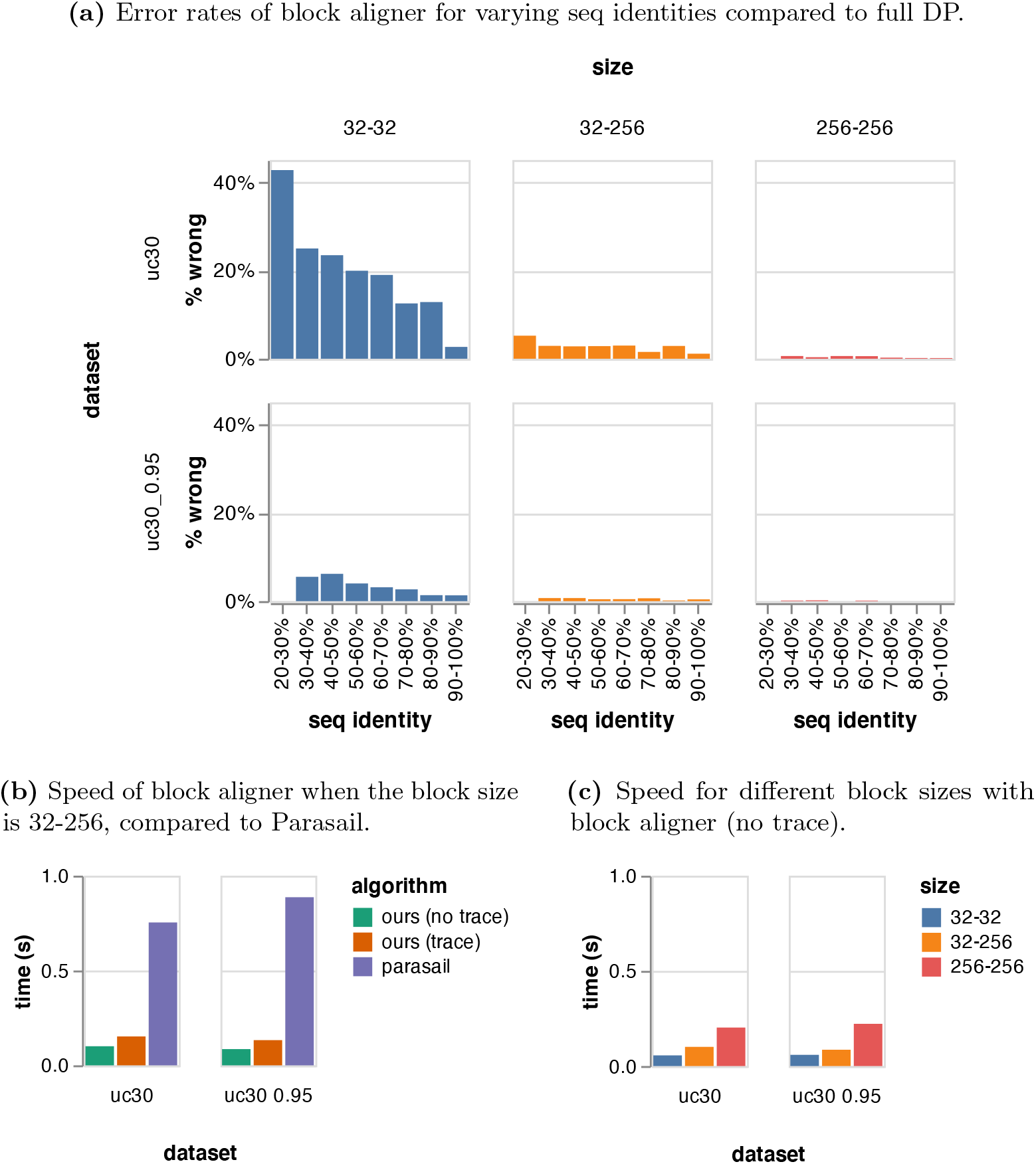
Global alignment of proteins from the Uniclust30 dataset, with block aligner.

In Figure 5a, we see that, as expected, the error rates increases as the sequence identity decreases. Additionally, with lower sequence coverage in the “uc30” dataset, the error rate of using a small block size becomes very high. For this data, it seems that a block size of 32-256 is a good balance between accuracy and speed, allowing it to be 7-9 times faster than Parasail [16], while maintaining similar accuracy. It is only a slightly slower than a fixed block size of 32 in Figures 5b and 5c, which means that the block growing heuristic works well at automatically staying on the fast path of not growing the block whenever possible. An important, yet subtle benefit of this is that we do not need to tune any parameters to handle proteins of wildly differing sequence identities, it just *works*. Additionally, block aligner scales sublinearly when the block size increases, due to the well-optimized SIMD DP and prefix scan code.

When the block size is 32-256, the average percent error of the wrong scores produced by block aligner is 13.7% for “uc30” and 4.4% for “uc30 0.95”. Compared to full DP on Uniclust30 proteins, on average only around 40% of the full DP matrix is computed with block aligner, when the block size is 32-256.

### Sequencing Reads Benchmark

Our first evaluation on real DNA data is comparing block aligner to full DP on global alignment of reads ranging from around 100 to more than 25,000 base pairs. The results are shown in Table 1.

**Table 1:** Global alignment of DNA reads with block aligner. The scores produced by block aligner are compared to full DP global alignment.

| dataset | block size | read pairs | wrong | error rate | wrong % error |
| --- | --- | --- | --- | --- | --- |
| illumina | 32-32 | 100000 | 0 | 0.0% | 0.0% |
| nanopore 1kbp | 32-64 | 12477 | 377 | 3.0% | 20.1% |
| nanopore 25kbp | 32-256 | 1734 | 100 | 5.8% | 8.2% |

We then compared our adaptive block shifting approach to static banding [13] and adaptive banding [14] in Figure 6. We use different length of prefixes of the ∼25kbp sequences because it is common to only align portions of sequences in real aligners, like with seed-and-extend approaches [4]. To reinforce the benefits of adaptive approaches over static banding, we test how varying static band widths compare against block aligner and report the results in Figure 6a. Even with orders of magnitude smaller block sizes than band widths, block aligner is able to be similarly accurate as static banding in many cases. In Figure 6b, Block aligner performs better than adaptive banding in many cases, which can likely be attributed to how adaptive banding only examines the center of the band for the max score. Also, the percentage of better scores decreases when we increase the length of the prefixes of the 25kbp Nanopore reads, which suggests that the suffix region of the sequences is much more challenging to align greedily. As expected, when block aligner is allowed to grow its block size from 32 to 64, it is more accurate.

**Figure 6:**
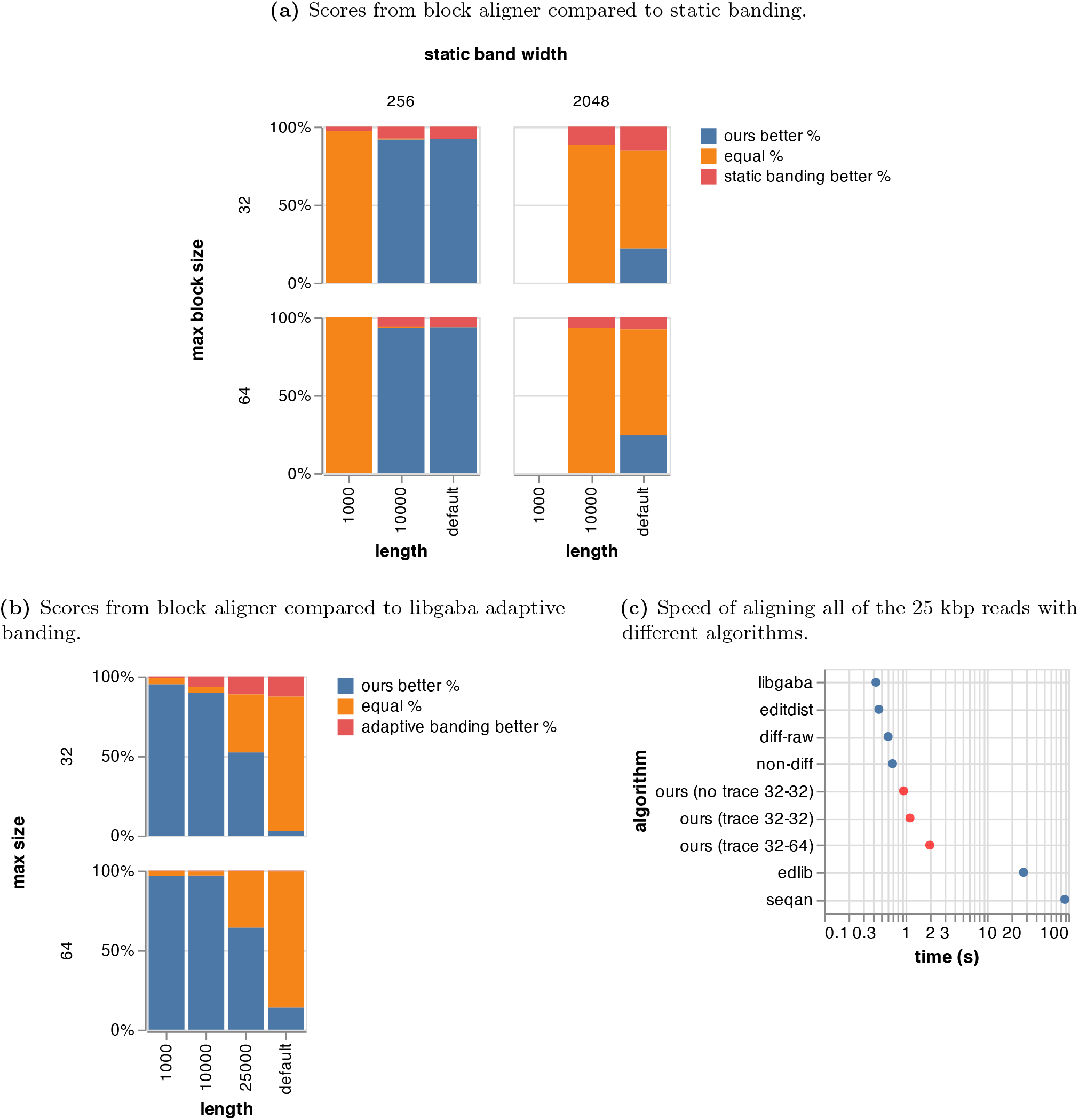
Comparison of block aligner with other algorithms on 25kbp Nanopore reads. For the comparisons, we look at the percentage of scores that were greater (better), because the compared algorithms would underestimate the score if it were wrong. The “default” length is the full length of the sequences (*∼*25 kbp). Other lengths indicate a prefix of the full sequence of that length is used. The *X*-drop threshold is 50 for all experiments except in (a) where it is *−∞*. In (c), traceback paths are not computed, but trace data is stored during alignment if possible.

In terms of speed, block aligner is around three times slower than the well-optimized libgaba [14]. This is likely due to 1. the vertical/horizontal vectors that require extra prefix scan steps to resolve dependencies, the wider 16-bit SIMD lanes in block aligner instead of 8-bit lanes with the difference recurrence method [14], and 3. a more complex block-based algorithm that handles growing and alternating between horizontal or vertical shifts.

## Discussion

Our results show that block aligner performs well in terms of speed and accuracy with a variety of different datasets. However, it is still advisable to use other algorithms in specific scenarios. For example, Wavefront Aligner [18] will likely perform better for computing affine-gap alignment with positive edit weights on sequences with high seq identities, due to its ability to quickly identify runs of exact matches. However, for general scoring methods, it is difficult to obtain the same speedups as Wavefront Aligner with the same accuracy guarantee.

We believe that block aligner fills the niche of efficiently aligning sequences of varying sequence identities with complex scoring matrices for larger alphabets. Before, variants of Farrar’s algorithm [15] has been commonly used for protein sequence comparisons [1, 16, 17], but now, block aligner is a strong contender for pairwise global alignment and X-drop alignment (*e*.*g*., after seeding) with its high speed and minimal accuracy loss. Of course, block aligner can also be used for aligning short or long DNA reads with large gaps using both negative and positive scores, similar to adaptive banding [13]. Block aligner can also be used for quickly obtaining a lower bound on the alignment score by purposefully constraining the block size, which may be useful in filtering sequences.

In terms of the implementation, there are a couple of limitations so far, including the accuracy/efficiency trade off with the step size. This problem arises because in practice, computing the shift direction and performing other checks between block shifts is slow, so we use a larger step size to spend more time on SIMD DP computation. However, this will slightly lower the accuracy compared to using a small step size like with adaptive banding [13]. We expect this to account for some of the differences in Figure 6b. Also, being a greedy algorithm, block aligner has trouble with low complexity or repeating regions, like with adaptive banding. These accuracy issues typically can be solved by using larger block sizes, though at the expense of speed.

## Conclusion

We propose a new algorithm, block aligner, for pairwise sequence alignment, and we show that it works well on real proteins and sequencing reads.

For future work, there is room to optimize the implementation for computing rectangular regions and how that interacts with the code for controlling block shifting and growing, to allow the step size to be smaller for more accuracy, without sacrificing speed. Additionally, more SIMD instruction sets, like AVX-512 and ARM Neon can be supported to allow greater portability and speedups on certain hardware. Also, position specific scoring matrices or additional alignment modes (*e*.*g*., local alignment) may be supported to expand the usefulness of block aligner in aligning proteins.

Sequence alignment is a core computational primitive in many bioinformatics workflows. Given this, it is important to explore the trade off between efficiency, flexibility, and accuracy of alignment algorithms. Block aligner is a practical step in this direction, and it reveals new methods for approaching the sequence alignment problem that may impact the broader bioinformatics community.

## Abbreviations

AVX: Advanced Vector Extensions
DP: Dynamic Programming
SIMD: Single Instruction Multiple Data

## Competing interests

The authors declare that they have no competing interests.

## Author’s contributions

DL designed the algorithm, created the implementation, performed experiments, and wrote the paper. MS gave feedback, provided experiment data, and wrote the paper.

## Acknowledgements

We thank Hajime Suzuki for helpfully answering and discussing many of our questions. Additionally, we acknowledge Twitter for connecting us together to make this paper possible.

## Author details

^1^University of California Los Angeles, 90095 Los Angeles, USA. ^2^Seoul National University, 1 Gwanak-ro, 08826 Seoul, Korea.

## Notes

https://github.com/Daniel-Liu-c0deb0t/block-aligner

### Competing Interest Statement

The authors have declared no competing interest.

